# How to count your bugs?

**DOI:** 10.1101/2021.02.08.430281

**Authors:** Ksenia S. Onufrieva, Alexey V. Onufriev

## Abstract

Ability to estimate local population density of an insect is critical in many fields, from pest management to conservation. No method currently exists that reliably connects trap catch with the insect population density, including the corresponding uncertainty. Here we report a simple and universal predictive relationship for a probability of catching an insect located a given distance away from the trap. This relationship allows to estimate, from a single catch, the most likely population density along with its statistical upper and lower bounds. To test the generality of this equation we used 10 distinct trapping data sets collected on insects from 5 different orders and major trapping methods: chemical-baited and light. For all of the datasets the equation faithfully 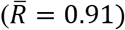 describes the relationship between location of an insect and probability to catch it. The ability to estimate absolute population density from a single trap catch will significantly improve our understanding of insect population dynamics and allow for more effective research, management, and conservation programs.

## Introduction

The question posed in the title of the paper may seem trivial: you catch your bugs with a net and count them. However, that procedure will tell you how many bugs are in the net and nothing abut what you really want to know – how many bugs are in the area around you.

Traps are crucial for monitoring insect activity and are widely used in pest detection and management programs (Abell et al., 2015; Elkinton & Cardé, 1981; Tobin et al., 2013; Tobin et al., 2004), for evaluating biodiversity and planning conservation efforts (Bossart & Carlton, 2002; Larsson, 2016; New, 1996), and in research efforts (Casado et al., 2014; Contarini et al., 2009; Elkinton & Carde, 1980; Tcheslavskaia et al., 2002; Tobin et al., 2009). Therefore, extensive research has been conducted to evaluate trap efficiency (Cardé et al., 2018; Elkinton & Childs, 1983; Ferracini et al., 2020; Irish et al., 2013; Jactel et al., 2019), to estimate range of attraction (Byers, 2008; Byers et al., 1989; Dufourd et al., 2013) and catch probability (Gage et al., 1990), and to better interpret trap catches and to relate them to the absolute population density (Bau & Cardé, 2016; Kirkpatrick et al., 2019; Kirkpatrick et al., 2018; Miller, 2020).

Any quantitative analysis of trap catches must include a crucial step – directly relating trap catches to the absolute population density of an insect. The absence of a reliable and universal (applicable to all insects) procedure for this crucial step hinders conservation, management and research programs making it difficult to interpret catches, provide recommendations, develop management tactics and evaluate treatment efficacies. The availability of statistically reliable estimates of the absolute population density would significantly improve existing conservation and management programs by allowing them to optimize efforts based on the goal, available resources and the efficacy of the previous efforts. In research programs, this would significantly improve interpretation of results and facilitate optimization of existing tactics and development of new ones.

In a recent study we analyzed a wealth of trap catch data collected for European gypsy moth *Lymantria dispar dispar* (L.) and derived a simple mathematical relationship between catch probability and distance to a pheromone-baited trap, which, in turn, allowed us to connect the actual trap catch with the most probably population density, along with statistical bounds on absolute gypsy moth population density (Onufrieva et al 2020). However, the key question remained unanswered: is there a general relationship of this type that might apply to all insects and trap types? In this paper we demonstrate the generality of this mathematical relationship using several species of insects from various orders, and two major trapping methods: chemical baited and light attraction.

Insects are the most diverse group of organisms and it is highly improbable that their behavior with respect to attractants could be described by a universal law. However, if such a law were to be found it could have significant impact on the entire field of entomology. This work is about finding and validating such a law.

## Methods

### Data collection

We searched the literature for data on insect catches in traps located at various distances from the insect release points that satisfied the following conditions: (1) converged catch (meaning that the value did not increase substantially with increased trapping time, as defined in (Onufrieva et al., 2020) was reported for at least 4 distances between a trap and a release point, (2) number of insects released at large distance was the same or larger than at short distances, (3) no zero catch data points were reported between non-zero points. This search yielded 9 data sets on brown marmorated stink bug (*Halyomorpha halys*) (Kirkpatrick et al., 2019), codling moth (*Cydia pomonella*) (Adams et al., 2017), European pine sawfly (*Neodiprion sertifer*) (Östrand & Anderbrant, 2003), spotted wing drosophila (*Drosophila suzukii*) (Kirkpatrick et al., 2018), Western corn rootworm (*Diabrotica virgifera*) (Wamsley et al., 2006), Douglas fir beetle (*Dendroctonus pseudotsugae*) (Dodds & Ross, 2002), Southern pine beetle (*Dendroctonus frontalis*) (Turchin & Odendaal, 1996), macro-moths in families Erebidae (Merckx & Slade, 2014) and Sphingidae (Beck & Linsenmair, 2006).

### Analysis

We used the predictive relationship for a probability of catching an insect (*spT_fer_(r)*) located at a distance *r* from the trap that was developed for gypsy moth (Onufrieva et al. 2020) to investigate if it could be applied to other insects.

For gypsy moth, a wealth of data points is available (Onufrieva et al., 2020), which allowed us to come up with the most robust protocol for fitting Equation 1 (see Results). Specifically, for gypsy moth, males were released at distances 0, 15, 25, 30, 45, 50, 60, 75, 80, 100, 150, 200, 250, 300, 500, 600, 900, 1000, 1200, and 1500 m from pheromone-baited traps, therefore, short and long distances were balanced and had equal weight in determination of *D_50_*. The gypsy moth data set is also unique in that 12 distinct points are available for large values of r (r > 75 m). The availability of multiple data points at long distances had previously allowed us (Onufrieva et al., 2020) to come up with what we believe is the most accurate estimate of *D_50_* = 26 ± 3 m, which was based on a log-log fit for long distance data points only. However, data available for the other insects studied here does not include *spT_fer_(0)* and experimental design is often unbalanced, including either mostly short or long distances, and not very many of them. To overcome this limitation, we developed a 2-step protocol for fitting Equation 1 to data that is missing *spT_fer_(0)*. Step 1: Use untransformed data to estimate *spT_fer_(0)* by fitting Equation 1 to the experimental data points (we employed JMP^®^ Pro 15, SAS Institute, 2019). Step 2: Use *spT_fer_(0)* from Step 1 in Equation 2 to estimate *D_50_* by fitting Equation 2 to the log-transformed experimental data points (JMP^®^ Pro 15, SAS Institute, 2019). This 2-step procedure ensures that the catches at large distances are given equal weight to the catches at short distances. For insect data that includes experimentally measured *spT_fer_(0)*, only step 2 should be used.

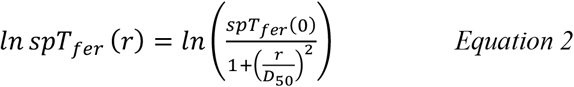

We tested this protocol for gypsy moth and estimated *spT_fer_(0)* = 0.15 and *D_50_* = 45 ± 5 m. This *spT_fer_(0)* is lower than the actual experimental *spT_fer_(0)=0.37* observed in the field (Onufrieva et al. 2020). Using the actual *spT_fer_(0)* in untransformed and log-transformed model yielded *D_50_* = 21.7 ± 3 m and *D_50_* = 27.3 ± 3 m, respectively. The latter value is closest to the one obtained previously, which supports the use of the 2-step fitting procedure including the log-transformed 2^nd^ step. The estimate of *D_50_* obtained using the 2-step protocol proposed for the datasets missing *spTfer(0)* is higher than the estimates obtained using the other two methods, but nevertheless Equation 1 with their respective parameter sets approximates the experimental data reasonably well (Fig. 1) in all three cases.

**Figure 1:**
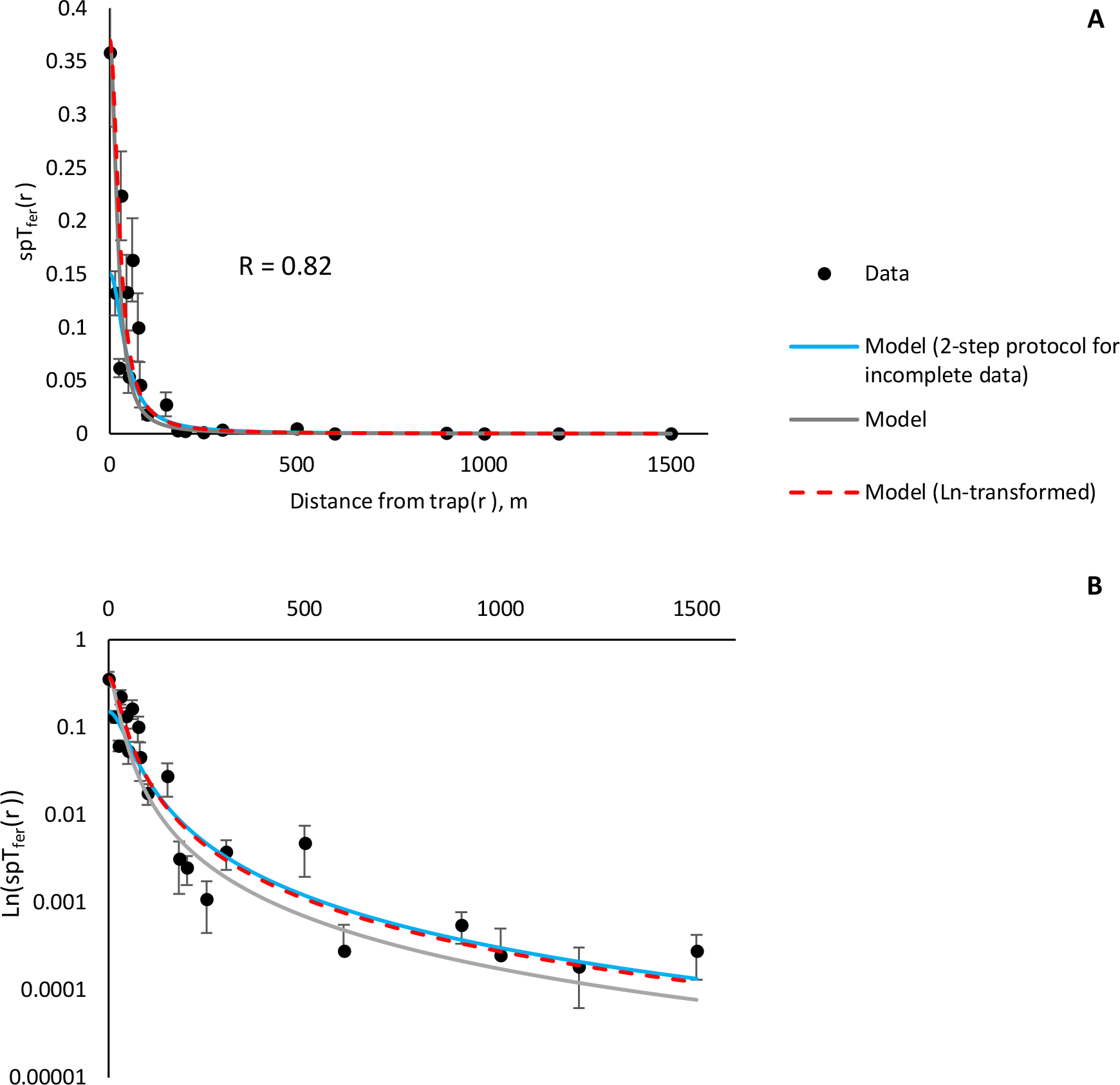
Proportion of gypsy moth males caught in pheromone-baited traps placed at various distances from the release point (±SEM). Error bar is not shown when smaller than the symbol size. Panel A, *spT_fer_(r)* vs. *r*, illustrates quality of the fit to Equation 1 at all distances; panel B, *ln(spT_fer_(r))* vs. *r*, illustrates the fit quality at large distances from the pheromone-baited trap.

Both probability of catch in the immediate proximity to the trap *spT_fer_(0)* and *D_50_* are crucial for establishing a relationship between catch probability and distance to a baited trap, deriving bounds on absolute population density, and estimating the most probable population density of an insect (Fig. 3).

**Figure 2:**
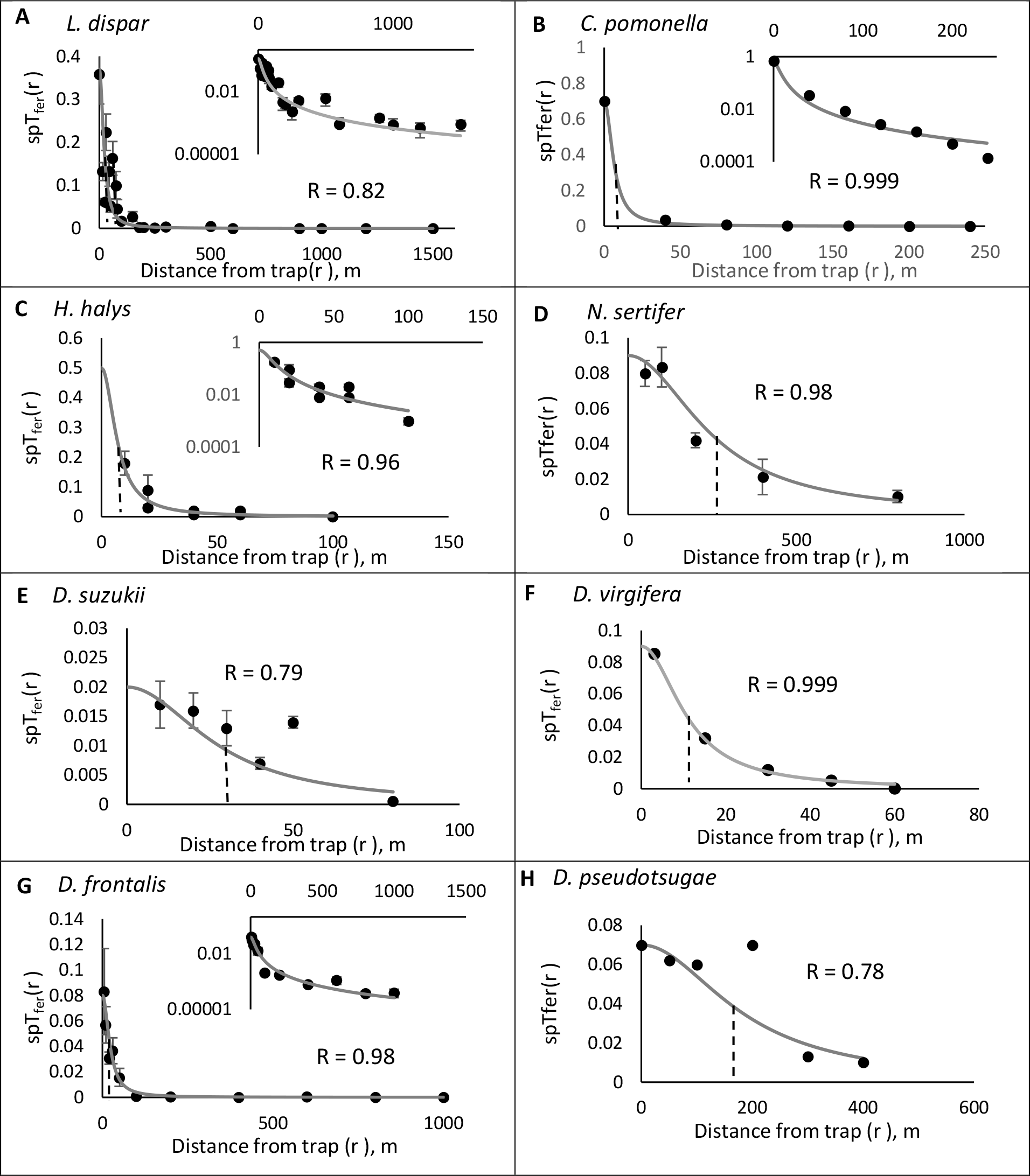
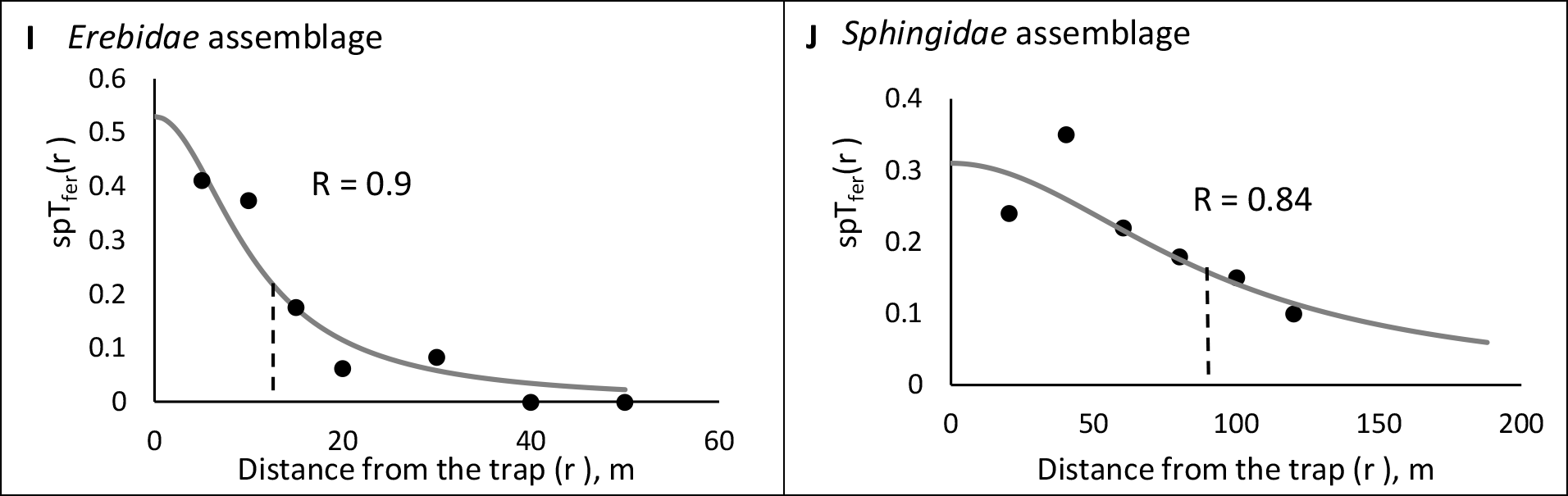
Proportion of insects caught in pheromone-baited traps placed at various distances from the release point (±SEM, where available). Black dots represent experimental data, grey line represent log-log model with *spT_fer_(0)* obtained using untransformed data. Black dashed lines mark *D_50_* estimated from the data as a distance where *spT_fer_(r)* = *1/2spT_fer_(0)*. In *C. pomonella* (A), *H. halys* (B) and *D. frontalis* (G), we included graphs to show fit at the large distances from the trap (ln(*spT_fer_(0)*) vs *r*), where trap catches are very low.

**Figure 3:**
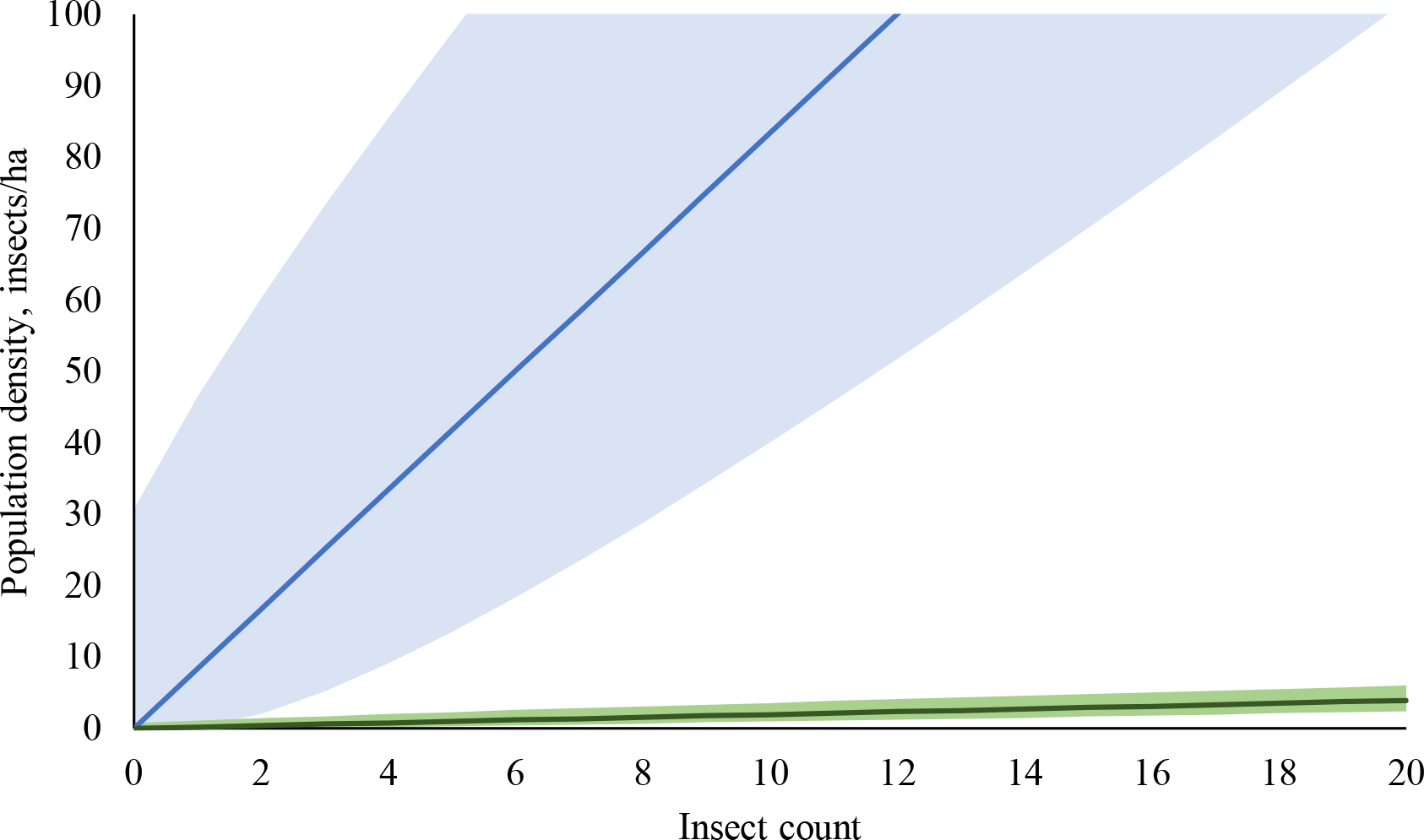
Absolute codling moth (blue) and European sawfly (green) population densities as a function of catch (insect count) in baited traps. Light blue and green areas indicate the ranges between lower and upper bounds with 95% probability for codling moth and European sawfly, respectively. Dark lines in the middle indicate the most probable average densities 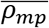. For the same number of insects caught, the corresponding population densities differ by nearly two orders of magnitude.

To derive bounds on average population density 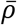, we use the procedure described in (Onufrieva et al 2020). Once *spT_fer_(0), D_50_* and *R_max_* are estimated, we define:

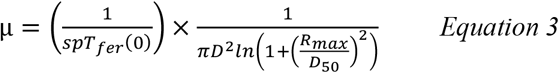

With that, the lower and upper bounds on the average density 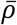

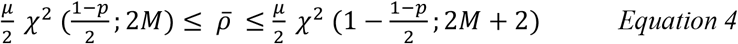

where M is the number of insects caught, p is the confidence level (p = 0.95 here) and *χ*^2^ (*q, n*) is the quantile function (corresponding to a lower tail area *q*) of the *χ*^2^ distribution with *n* degrees of freedom (Table 2).

The most probable average male density in the trapping area is:

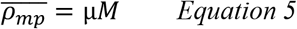

To convert the male density to number of males per ha, and assuming *D_50_* and *R_max_* are given in meters, μ in Equations (4) and (5) needs to be multiplied by 10,000.

We note that the probability of catching an insect located in the immediate proximity to the trap *spT_fer_(0)* provides a reference point for the rest of the trap catches, which is why it is important to measure *spT_fer_(0)* empirically, since as we saw in the example with gypsy moth data, estimating *spT_fer_(0)* by fitting Equation 1 to the experimental data is possible, but may not always match the experimentally obtained *spT_fer_(0)*, which, in turn, may lead to an over- or underestimated *D_50_*. In western corn rootworm (see Results), our estimated *D_50_* = 11 m agrees with results reported by Wamsley et al. (Wamsley et al., 2006), who observed significant drop of trap catches beyond 30 m away from the trap. However, trap catch collected at the distance of 16 m away from the trap is also significantly lower compared to the catch in a trap located 3 m away (Fig. 2E). This, once again, demonstrates the importance of measuring *spT_fer_(0)* empirically rather than estimating it by fitting Equation 1 to an incomplete experimental dataset. In Douglas fir beetle, *D*. pseudotsugae, previous studies reported that traps attracted beetles from at least 200 m (Dodds & Ross, 2002), but beyond this distance the recapture rate drops, which agrees with our estimate of *D_50_* =184 ± 33 m (see Results).

## Results

The main result of this work is the universal predictive relationship for a probability of catching an insect (*spT_fer_(r)*) located at a distance *r* from the trap.

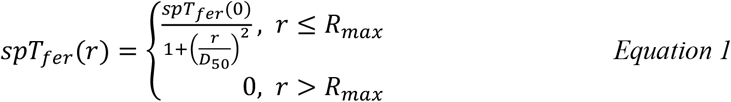

where *spT_fer_(0)* (Miller et al., 2010) is the probability of catching an insect located in the immediate proximity to a baited trap and *D_50_* is the distance from a baited trap at which the probability to catch an insect is ½ of the probability to catch an insect in the immediate proximity to the trap (*spT_fer_(0)*).

Results of the analysis conducted to estimate *spT_fer_(0)* and *D_50_* for the studied insects are shown in Table 1 and Fig. 2. In all cases, the estimates of *D_50_* obtained using untransformed and log-transformed data were very similar, within standard error of the mean (Table 1). The log-transformed values correspond to the 2-step protocol described in Methods, while only the 1^st^ step was used to fit the untransformed data to obtain the corresponding *spT_fer_(0)* and D_50_. The similarity of D_50_ values obtained using two different fit procedures further supports robustness of Equation 1.

**Table 1:** Estimates of probability to catch an insect released in the immediate proximity to the trap (*spT_fer_(0)*) and *D_50_* for various insects in orders Lepidoptera, Coleoptera, Hymenoptera, Diptera, and Hemiptera. Experimental R_max_ is listed, except for Sphingidae (marked *), which was estimated using method described in Miller et al. (2015).

| Insect | Trap type | $spT_{fer}(0)$ | $D_{50} \pm \text{SEM}$, m<br>untransformed data | $D_{50} \pm \text{SEM}$, m<br>**log-transformed**<br>**data** | $R_{max}$, m |
| --- | --- | --- | --- | --- | --- |
| Gypsy moth<br>(*L. dispar*) | Pheromone | 0.37 | $21.7 \pm 3$ | **$27.3 \pm 3$** | 1600 |
| Codling moth<br>(*C. pomonella*) | Pheromone | 0.7 | $9 \pm 0.1$ | **$6.5 \pm 1$** | 260 |
| Brown marmorated<br>stink bug (*H. halys*) | Pheromone | 0.5 | $7 \pm 3.5$ | **$7 \pm 0.9$** | 130 |
| European pine<br>sawfly (*N. sertifer*) | Pheromone | 0.09 | $216 \pm 39$ | **$250 \pm 21$** | 1040 |
| Spotted wing<br>drosophila<br>(*D. suzukii*) | Chemical | 0.02 | $45.4 \pm 16$ | **$27.7 \pm 7$** | 90 |
| Western corn<br>rootworm (*D.*<br>*virgifera*) | Pheromone | 0.09 | $11 \pm 0.2$ | **$11 \pm 0.4$** | 60 |
| Southern pine beetle<br>(*D. frontalis*) | Pheromone | 0.08 | $19.4 \pm 2.8$ | **$23 \pm 2.8$** | 1000 |
| Douglas fir beetle<br>(*D. pseudotsugae*) | Pheromone | 0.07 | $251.5 \pm 115$ | **$184 \pm 33$** | 600 |
| Erebidae assemblage | Light | 0.52 | $11.6 \pm 3$ | **$10.6 \pm 1.4$** | 40 |
| Sphingidae<br>assemblage | Light | 0.31 | $97 \pm 27$ | **$91.6 \pm 8$** | 175* |

Once parameters that enter Equation 1 have been obtained, one can relate, quantitatively via Equations 3, 4 and 5, the number of trapped insects with the actual population density, as exemplified for two insects in Fig. 3.

## Discussion

The key result of this work is the demonstration that a universal equation exists that faithfully describes the relationship between the probability to catch an insect and how far it is from the trap. The relationship is a simple formula with only 2 key parameters: *spT_fer_(0)*, which is a probability to catch an insect released in the immediate proximity to the trap and *D_50_* which we define as the distance from a baited trap at which the probability to catch an insect is ½ of the probability to catch an insect released in the immediate proximity to the trap (*spT_fer_(0)*). The strength of this definition is threefold: (1) it directly corresponds to what can be measured in field experiments and, (2) the concept of *D_50_* can be easily illustrated on the graph of *spT_fer_(r)* vs. *r*, from which *D_50_* value can be immediately estimated, at least approximately, as the value of *r* at which *spT_fer_(r)* = *1/2spT_fer_(0)* (Fig. 2), and (3) the definition applies to any trap type.

To understand the biological meaning of *D_50_*, and its possible relationship to insect physiology, we compared D_50_ values derived from the trapping experiments with direct measurements of insect physiological response to appropriate attractant, where available. In gypsy moth, we estimated *D_50_* = 26 ± 3m (Table 1), while Elkinton et al. (Elkinton & Cardé, 1984) observed wing fanning starting at a distance of 20 m from the pheromone source. Our estimate of *D_50_* for European sawfly (*D_50_* = 250 ± 21 m) agrees with the results of behavioral studies reported by Östrand et al. (2000), who observed response in *N. sertifer* to pheromone sources located 200 m away. Based on the agreement of our results with physiological studies, we suggest that qualitative biological meaning of *D_50_* is effective attractive distance at which the probability that the lure elicits a response from is substantial. To formulate a more quantitative relationship will require more detailed physiological experiments than currently available.

In gypsy moth, numerical values of *D_50_* and plume reach described by Miller et al. (2015) happen to be similar, but the match is purely coincidental and does not hold for most insects studied here. For most insects the values of plume reach (a pheromone-specific concept), and *D_50_* (a universal characteristic of any trap) differ significantly.

One of the most striking results of this study is that the same number of insects caught in a trap may translate into order of magnitude different population densities in the field (Fig. 3). The qualitative explanation is that the population density is sensitive to parameters of the trap-insect system, particularly *D_50_*, and *spT_fer_(0)*. Without knowing these key characteristics, based on the trap catch alone, one cannot make any quantitative assessment of what the actual insect population might be. The meaning of “catch zero” and “catch one” become clear only in light of the established relationship with the statistical bounds on the population density. When no insects are caught in the trap, we can conclude that, even though the insects might still be present in the field, their population density cannot exceed the specific threshold (upper bound, 95% confidence, Fig. 3). Likewise, if only a single insect has met its sad end in the trap, one can conclude that the actual population density cannot, with 95% confidence, be lower than the appropriate lower bound (Fig. 3).

It is remarkable that the simple Equation 1 works so well (average R = 0.91) across 5 orders of insects collected using very different attractants, such as chemical and light, selected randomly from the literature based on the available data despite the fact that parameters of analyzed trap-insect models vary widely: *D_50_* ranged 6.5 – 250 m and the estimated probability of catch in the immediate proximity to the trap *spT_fer_(0)* ranged 0.02 – 0.7 (Table 1). This universality is the consequence of the universal set of principles that we applied to trapping of all insects: two-dimensional active movement space (insects following the terrain), finite active life span, and converged trap catches (collection time is long enough) used in well-designed trapping experiments.

Importance of conservation and pest management programs cannot be overstated as climate change, loss of biodiversity, and biological invasions remain the most serious environmental problems facing society. Inability to interpret insect trap catch data quantitatively, which includes directly relating trap catches to the absolute population density of an insect, hinders conservation, management, and research programs by making it difficult to provide recommendations, develop management tactics and evaluate treatment efficacies. The universal method reported here fills a key knowledge gap: it allows rigorous estimation of the most likely insect population density, along with the corresponding upper and lower bounds, from the number of insects caught by a single trap. The method is universal, in that it can be used for any trap-insect system. We believe this method will help develop technologies for improved insect population detection and management, but most importantly, will help drive future basic and applied research in multiple areas of entomology and ecology.

## Acknowledgements

We thank Dr. Jim Miller for help with locating relevant experimental work and for many stimulating discussions. We thank Dr. Andrew Liebhold for useful comments.

## Appendix

**Table 1:** Quantile function of the *χ*^2^ distribution with n degrees of freedom, p = 0.95, to be used in *Equation 4*.

| # males caught | $\chi^2 ((1 - p)/2, 2M)$ | $\chi^2 (1 - (1 - p) / 2, 2M+2)$ |
| --- | --- | --- |
| 0 | 0 | 7.37776 |
| 1 | 0.0506356 | 11.1433 |
| 2 | 0.484419 | 14.4494 |
| 3 | 1.23734 | 17.5345 |
| 4 | 2.17973 | 20.4832 |
| 5 | 3.24697 | 23.3367 |
| 6 | 4.40379 | 26.1189 |
| 7 | 5.62873 | 28.8454 |
| 8 | 6.90766 | 31.5264 |
| 9 | 8.23075 | 34.1696 |
| 10 | 9.59078 | 36.7807 |
| 11 | 10.9823 | 39.3641 |
| 12 | 12.4012 | 41.9232 |
| 13 | 13.8439 | 44.4608 |
| 14 | 15.3079 | 46.9792 |
| 15 | 16.7908 | 49.4804 |
| 16 | 18.2908 | 51.966 |
| 17 | 19.8063 | 54.4373 |
| 18 | 21.3359 | 56.8955 |
| 19 | 22.8785 | 59.3417 |
| 20 | 24.433 | 61.7768 |
| 21 | 25.9987 | 64.2015 |
| 22 | 27.5746 | 66.6165 |
| 23 | 29.1601 | 69.0226 |
| 24 | 30.7545 | 71.4202 |
| 25 | 32.3574 | 73.8099 |
| 26 | 33.9681 | 76.192 |
| 27 | 35.5863 | 78.5672 |
| 28 | 37.2116 | 80.9356 |
| 29 | 38.8435 | 83.2977 |
| 30 | 40.4817 | 85.6537 |

